# The dynamic face of rarity: tracking species at risk beyond space

**DOI:** 10.64898/2026.10.11.758290

**Authors:** Everton A. Maciel

## Abstract

Avoid extinction of rare species is critical for biodiversity conservation. Rabinowitz proposes rare species is pattern that emerge from combination of three parameters: geographical range, habitat specificity and population size. Several papers have been looked to improves this method over last years, but they often neglect the temporal dimension, crucial for understanding the extinction process. This study introduces a dynamic framework that model the three dimensions of rarity as a continuous three-dimensional space, integrating vector-based representations and an ecological vector field to quantify rarity, temporal dynamics, displacement and equilibrium. Derivative equations are used to describe the temporal evolution in that system which is integrate to account the shift in that space. This framework allows to reinterpret ecological change as a trajectory though space, where each axis corresponds to an ecological dimension. This multidimensional representation distinguishes intrinsic rarity from extinction risk and can identify demographic declines, habitat restriction, or range contraction. The method can be applied to a single species or a set of species, using short or long sampling time intervals. A test across freshwater fish, birds, amphibians, and terrestrial plants showed that rarity dimensions can change independently among species. By monitoring the shifts in this dimension simultaneity, we can plan the conservation by priority species that are shaking their dimensions over time. Unlike static assessments that already exist, this method provides a nuanced understanding of a rare species’ vulnerability as a dynamic representation of temporal changes. An R package and application guide are provided, helping conservation practitioners use this approach.

## 1. Introduction

Avoid extinction of rare species is critical for biodiversity conservation. In this context a growing body of literature has acknowledged that species with a narrow geographical range, habitat specificity, and small population size are prone to extinction (Davies et al., 2004; Harnik et al., 2012; Maciel et al., 2023; Matthies et al., 2004; Wang et al., 2018). The local extinction rate is believed to be inversely proportional to the population size (Brown, 1986) because fitness decreases in species with small populations due to genetic drift and inbreeding (Nabutanyi and Wittmann, 2021), as well as the difficulty of finding mating partners (Cuddington et al., 2015). Additionally, habitat specificity has been linked to increased vulnerability to extirpation (Reinhardt et al., 2005; Stöcklin and Fischer, 1999), because certain plant species that specialise in a particular habitat may have higher local extinction rates due to lower seed longevity compared to generalist species (Stöcklin and Fischer, 1999). This may be attributed to the fact that habitat specialist species are more likely to have their habitat destroyed than generalist species (Marini et al., 2010; Reinhardt et al., 2005).

Rabinowitz proposed that rarity is a pattern that results from the interaction of three parameters: population size, habitat specificity, and geographical range (Rabinowitz, 1981). Over the past three decades, a dozen studies have proposed novel indices based on the Rabinowitz scheme. Some of those indices adhered more closely to Rabinowitz and restricted themselves to the parameters She recommended, like a combination of species specific to the habitat and restricted to a single plot (Acosta et al., 2009). Other index took into account a greater number of variables, such as habitat, altitude, and phytogeographical distribution on a regional and global scale (Betanio and Buenavista, 2018), or a combination of abundance, number of locations, islands and natural reserves where a species occurred (Borges et al., 2000)Rabinowitz’s parameters were integrated with different stages of species development to create composite rarity indices (Hodgson, 1993). Others researcher used one parameter of Rabinowitz to weights to rare species that exist below a specified occurrence criterion (Leroy et al., 2013). Since most of the previously proposed indices did not account for all three of Rabinowitz’s dimensions at the same time, the inverse of the sum of Rabinowitz’s three dimensions was also proposed somewhere (Maciel, 2021). More than mere competition, the wave of index development demonstrates progress in this field. However, they sought to improve Rabinowitz’s methods solely in spatial terms, overlooking the interaction between space and time.

The temporal and spatial dynamics of biodiversity loss pose significant challenges for conservation and management strategies (Brown et al., 2019; Diniz et al., 2021; Kalaba et al., 2013; Qu et al., 2022). Given the critical role of the time and space iteration on the evolution (Levin, 2004) and extinction of species (Cobben et al., 2015), a comprehensive conservation approach should consider both spatial and temporal scales (Ahmadi and Sedghamiz, 2007; Nordén et al., 2014). These is important because while the spatial patterns of biodiversity informs where conservation actions are needed, the temporal patterns inform the urgency in which such strategies should be implemented (Pressey and Taffs, 2001). Moreover, given the limited financial resources available for biodiversity conservation (Hemming et al., 2022; Holdgate, 1992), identifying where and how fast the actions should be taken to conserve species can help to allocate resources more equitably and efficient. Methods in conservation for instance have to be implemented in the context of the time and space.

Some authors have proposed that the temporal shifts in the distribution of taxa or entire communities can be investigated through the equilibrium principle, a fundamental assumption that underlying many ecological models and theories (Kean and Barlow, 2004; MacArthur and Wilson, 1963; Rasztovits et al., 2014). This framework posits that, depending on the spatial and temporal scale of observation, communities may be either in equilibrium or not (De Angelis and Waterhouse, 1987; Vetter, 2005). In this context, a community or population is considered to be in equilibrium when the number of individuals or the spatial distribution of species remains stable over time, as losses due to extinction are balanced by the continuous arrival or colonisation of individuals (Kuussaari et al., 2009). By contrast, non-equilibrium emerge from endogenous fluctuations operating across multiple levels of biological organisation, ranging from single populations and small sets of interacting species to entire ecological communities (Roy et al., 2020).

A variety of approaches can be used to assess equilibrium versus non-equilibrium dynamics in taxa or communities, often focusing on shifts in species populations or entire community (Law et al., 2019). These dynamics may be further influenced by external stressors such as climate change, habitat fragmentation, and other anthropogenic pressures. Species responses to such stressors vary considerably, reflecting differences in their ability to perceive environmental heterogeneity, as well as in their dispersal rates and persistence following disturbance (Liira et al., 2014). In this context, generalist species are typically less sensitive to environmental change due to their broader niche breadth and ability to persist in small or fragmented patches, whereas specialist species tend to be more vulnerable under similar conditions (Raatikainen et al., 2018). Moreover, common are better competitors that tend to outperform their rare counterparts when resources are scarce (Rabinowitz et al., 1984). Since species’ response to anthropogenic stressors differ, an intriguing question in this regard is how —and to what extent—species are expanding or retracting their ranges, populations, and habitat occupancy.

The Rabinowitz seminal paper established a cut-off value to put rare and common species apart (Rabinowitz, 1981). This cut-off values are the base of the methods and are applied to each of the three Rabinowitz’ dimensions (Maciel and Arlé, 2020). For geographical range dimension, the cut-off value is determined by the number of areas or latitude and longitude that a species is detected relative to entire area. Species that occur in 10% (cut-off value) or less of the sampled areas are considered geographically restricted, while species above that are widespread (Liu et al., 2020; Sætersdal and Birks, 1997). The habitat dimension considers species that are restricted to one habitat type (cut-off value) as habitat specialists (Söderström et al., 2007), while species that are above this are habitat generalist. For population size, species with two or less individuals per sampled unit (cut-off value) are considered to have a small population size while above this they have large population size (Pitman et al., 1999). This cut-off value has been the base of the studies of the thousands of studies attempted to conserve rare species. However, non-studies were made in attempted to implement this cut-off value in the space time dimension.

This paper uses the principle of equilibrium to propose a method to measure the dynamic of rarity of species take into account the space and time to quantify the shift of a taxa based on the three Rabinowitz dimensions. Rather than treating Rabinowitz’s parameters, geographical range, habitat specificity, and population size, as static or independent metrics as it was done before (Maciel, 2021) this paper defines them as coupled state variables evolving jointly over time in a continuous three-dimensional Cartesian space (Dorier, 1995). Using vector calculus and differential equations, this formalise the instantaneous and cumulative temporal dynamics, system equilibrium conditions, vector metrics that quantify both the magnitude and directional trajectory of species rarity across ecological communities (Carcassi and Aidala, 2023). This approach captures the ecological reality that abundance, spatial distribution, and habitat occupancy dynamically shift across temporal and spatial scales (Belmont et al., 2022; Mao and Colwell, 2005). The spatial shift of any taxon is measure through the shift in geographical range, the habitat occurrence and population size over time. The equilibrium could be achieved if one or all Rabinowitz dimensions of a taxa remain constant over time, while the non-equilibrium results for the increase or decrease of these dimensions over time. First, (1) the Rabinowitz parameters are integrated to vector principle to create a theoretical three-dimensional space (Maciel, 2021), where each taxon is conceptually considered as a point in three dimensions space (Brown and Maurer, 1987). Then, (2) the physical vectorial principle is used to create on equation to measure the rarity dynamic of the Rabinowitz dimensions base on the interactions of the space and time. Finally, (3) the occurrence of species is used to calculate the equilibrium of species over time and space.

## 2. Defining a 3D vectorial space of Rabinowitz

### 2.1. Definition of the 3D Rabinowitz state space

From this point, we will assume that each Rabinowitz’ dimension is a positive integer that is representative of a start point (*A*), and an endpoint (*B*) defines the cut-off value established by Rabinowitz. This cut-off value essentially creates a space limit, enabling the division of species into rare and common categories. Each dimension can be visualised as a line segment (*AB*) connecting these two points. We can represent the position of a given rarity dimension using an oriented segment between the origin *(0, 0, 0*) and the extremity (*a, b, c*) of these segments. The segment indicates the range of values that a taxon can have in that rarity dimension. To quantify the length of the oriented segment (*AB*), we can use a versor *u*.

Now let’s define Rabinowitz variable not as independent dimension as it was done before (Maciel, 2021) but as coupled state variables evolving jointly in time. The versor u represents the non-negative real number that corresponds to the rarity value of a taxa in a given rarity dimension. It indicates how far the taxon is from the origin in that dimension, with larger values representing higher rarity. We then have another versor u whose origin is (*0, 0, 0*) and extremity is the ordered term (*a, b, c*) in the three-dimensional Rabinowitz space (*ℝ*^3^). This versor represents the specific rarity combination of a taxa across all three dimensions geographical range (*GR*), habitat specificity (*HS*), and population size (*PS*). By using versors to show and measure the position and length of the directed segments, we can represent and study how rare different taxa are in the three-dimensional Rabinowitz space. This implies that Rabinowitz dimensions are part of a function or a set of functions that belong to a real variable *x*, with values of *f(x)*, that are defined as intervals of *ℝ*^3^, means that all values in x are part of these intervals, which could be defined as:

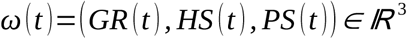

where *ω* (*t*) belong to a *ℝ*^3^, *GR(t)* represents the geographical range occupied by a taxon at time, *HS(t)* represents habitat specificity (or conversely, habitat breadth), and *PS(t)* represents population size, which together compose the *ℝ*^3^ space.

### 2.2. System dynamics and vector field

From now on we will assume that the joint temporal trajectory of the state variables is governed by an underlying ecological vector field *F* : *ℝ*^3^ ⟶ *ℝ*^3^ :

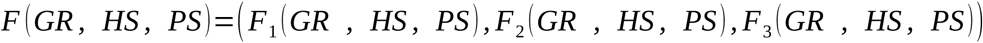

where *F* is associate each point in the space to a vector *F*_1_ *, F*_2_ *, F*_3_. The instantaneous rate of change of the rarity-state vector is given by the system of differential equations:

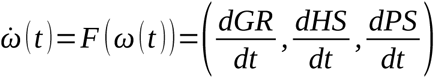

where *ω̇*(*t*) represents the rate of change of the rarity-state vector through time and *F* (*ω* (*t*)) is a field evaluate the position of the current point. The terms 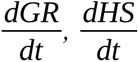 and 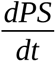 correspond to the rates of change of geographical range, habitat specificity, and population size, respectively. Together, these derivatives describe the instantaneous variation of a species within the Rabinowitz state space. Therefore, if we consider a time interval [*t*_1_ *, t*_2_], where t1 and t2 are, respectively, the final and initial instants where *t*_2_>*t*_1_, we can integrate the vector over time. Thus, the evolution of the system is given by:

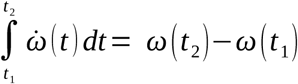

where *∫* is the integral of time, *`ω`* is a vector velocity, dt is an element of infinitesimal of the time, resultant of difference of *ω* (*t*_2_)*−ω* (*t*_1_). This means that the sum of all instantaneous variations over time results in the total variation of the system. Thus, each Rabinowitz dimension would correspond to the following system components:

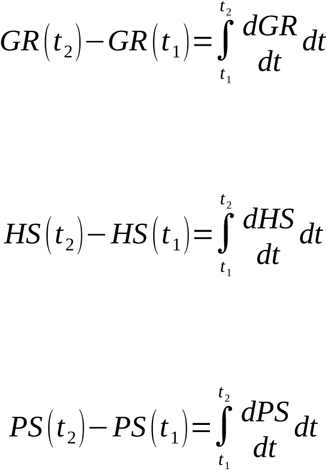

### 2.3. Cumulative rarity change and equilibrium conditions

To quantify the cumulative rate of aggregate variation across all three dimensions, we define the scalar metric d as the mean integrated change over time:

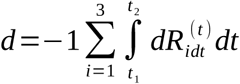

where *d* represent the cumulative of rarity change. The letter sigma ∑ is a summation of the elements of the three dimension that follow. Ri represents the *-th* Rabinowitz dimension (*R_1_ = GR, R_2_ = HS and R_3_ = PS*). At equilibrium, dimensional expansions and contractions cancel each other out, resulting in zero net shift in the taxon’s rarity position over time.

### 2.4. Ecological state of the variables

The empirical rate shifts for each dimension are calculated between initial (*t_1_*) and final (*t_2_*) sampling periods (*Δ_t_ = t_2_ – t_1_*). Geographical range (*GR*) is a factor which undermine the abiotic factors influences in species distribution limits on continental or global scale mostly (Brown et al., 1996). Therefore, the geographical range is determined by the presence of a taxa in a specific environment interval or across its entire dispersal range. This parameter can be measured using the geographical coordinates of a species (Maciel, 2021), the number of latitudinal bands where a species is found (Caiafa and Martins, 2010), or the area detected relative to the total sample area (Rey Benayas et al., 1999).

The GR is calculated based on the latitude and longitude of the species’ occurrences at the location where the data were collected. The gr for both *gr_i_* and *gr_f_* is calculated by the product of the latitude (*lat*) and longitude (*long*) ranges for each taxon. The shift in geographical range is give by:

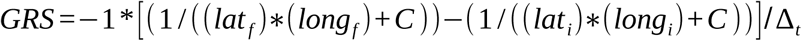

where the *GRS* is displacement of geographical range resulting from the difference between the final (*gr_f_*) and initial (*gr_i_*) geographical range of a taxon. C is a constant value added to the equation, potentially accounting for a baseline geographical range even at zero latitude and longitude. The *Δ_t_* is time interval resulting from the difference between the final (*t_i_*) and initial (*t_f_*) time, when the *gr_i_* and *gr_f_* were sampled. The difference between these gradients, scaled by the inverse of the time difference (Δt), gives you the overall change in geographical range.

Habitat specificity is a measure to indicates how specialised a species is in its habitat, meaning it either occurs in only one specific habitat or in multiple habitats. The habitat number refers to the instances of a species being present in a particular habitat. Initially, habitat was defined as a vegetation type (Kruckeberg and Rabinowitz, 1985), but more recent approaches have introduced various methods to quantify it, such as ecoregions (Kricsfalusy and Trevisan, 2014; Maciel and Martins, 2021) soil types for terrestrial organisms (Sperduto and Congalton, 1996), and different zones in the aquatic environment (Winter et al., 2002). The habitat number can represent any non-numeric parameter that characterises the habitat type of a species, such as soil type or vegetation type in the sampled area.

The habitat specificity shift (*HSS*) is calculated by the maximum number of habitats that a species can occur, and it is calculated by dividing the variation in habitats specificity (*Δ_hs_*) that a species exhibited by the time interval (*Δ_t_*) it took to varies.

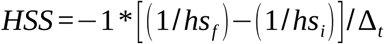

where *HSS* is the number of habitats resulting from the difference in the total number of at the final time (*h s_f_*) and the initial time (*h s_i_*). The *Δ_t_* is time interval resulting from the difference between the final (*t_i_*) and initial (*t_f_*) time. where *HSS* represents the change in the inverse of habitat shift (stability or persistence) at time *t*. *Δ_h s_* the change in the habitat specificity with respect to the state variable *S* (habitat shift). The equation incorporates these gradients through a subtraction, highlighting the change in stability between the initial and final states. Time dependence is factored in through the time difference (*Δt*), normalising the stability change over the observed time-frame.

The population size is a parameter that indicates the presence of a specific species in a particular area. The most commonly used measure for assessing local population size is the count of individuals (Maciel, 2021; Stien et al., 2022), but other measures such as individual density (Maciel and Guilherme, 2022), reproductive phenology (Duffy et al., 2013; Patrick et al., 2018), biomass (Dobson, 1995) and occupied territory count (Hoy et al., 2015) have also been utilized. In this context, population size can be any numerical value that represents the population of a species, such as density or the number of individuals within a specific sampling area.

The population size shift (*PSS*) is calculated by the maximum population size that a species presents in a given area, and it is calculated by dividing the population size variation (*Δ_ps_*) that a species exhibited by the time interval (*Δ_t_*) it took to varies. The magnitude of the value reflects the rate of change relative to the time interval. This allows direct comparison across different populations and time intervals.

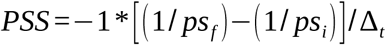

where *PSS* is the population size of a specie resulting from the difference in the number maximum population size (*ps*) at the final time (*ps_f_*) and the initial time (*ps_i_*). The *Δ_t_* is time interval resulting from the difference between the final (*t_i_*) and initial (*t_f_*) time.

## 3. Workflow for calculate rarity dynamic

### 3.1. Methods implementation

First, it uses the ‘as.character’ function to extract unique taxon names from the input data (Table 1) in order to create a species list. It then uses the range function to get the maximum and minimum latitudes and longitudes for every taxon. It calculates the geographical latitude and longitude ranges by utilising the diff function. The latitude and longitude ranges are then multiplied together and a constant of 1 is applied to get the geographical range of each taxon. Additionally, the habitat specificity is assessed by identifying the maximum number of habitats in which a taxon occurs, utilising the unique and length functions. Similarly, the population size is determined based on the maximum number of habitats in which a taxon is found, employing the max function. Any value that already exists in the same matrix by species is used to fill in any missing values in the data matrix. Put differently, a species that is found in four different habitats and whose habitat record indicates NA will have its NA substituted by any of the habitats in which it is found. These computations are performed for both time 1 and time 2 data. The results from the initial and subsequent surveys are merged into a unified matrix, containing the geographical range (*GR*), habitat specificity (*HS*), and population size (*PS*) for both time points. To standardise the values, the inverse of 1 is applied to *GR, HS*, and *PS*, yielding respective values ranging from 0 to 1 for each time period.

Subsequently, the differences in *GR, HS*, and *PS* between time 2 and time 1 are calculated, resulting in the displacement of geographical range, habitat specificity, and population size for each taxon. These displacements are then divided by the time interval (final time – initial time), yielding the geographical range shift (*GRS*), habitat specificity shift (*HSS*), and population size shift (*HSS*) for each taxon. Finally, the rarity dynamics (*rrD*) for each taxon are computed as the mean of the sums of *GRS, HSS*, and *PSS*. These values are subsequently multiplied by −1 to reverse the signs, ensuring that negative values indicate a decrease in parameters at time 2 relative to time 1, while positive values denote an increase. Values closer to 1 signify a more substantial increase or decrease in rarity dynamics.

### 3.2 Input data

The function was tested using real-world data from different sources and with different spatial and taxonomic resolutions. All datasets had to be organised according to the same format, containing: (1) species names in the first column, followed by (2) geographical coordinates, (3) habitat, and (4) population size for times one and two. Times one and two correspond to two surveys, which may range from hours to years, depending on the organism under study. Similarly, geographical coordinates can be recorded at a local scale (e.g. the latitude and longitude of plots) or at a continental scale (e.g. the centroid of grid cells). The only requirement is that the input data be in numerical format; for coordinates, these must be provided in decimal degrees. Habitat must be a character variable, which may represent temperature, elevation, vegetation type, and so forth. Population size must also be provided in numerical format, but it is otherwise fully flexible and can represent any attribute that reflects the population size of a species, such as density, basal area, number of individuals, etc. Observations must be provided for times one and two, and missing parameters should be left blank.

The first and second dataset consists of local-scale population dynamics data for one owl species and one salamander species, downloaded from the Government of Canada website. The owl data consist of occupancy records from ten stations across four time series (2002– 2003, 2003–2005, 2003–2011, and 2011–2012). The coordinates are provided in UTM, the habitat consists of four vegetation types (North Mountain/North Aspy Valley, Jigging Cove/South Mountain, MaryAnn Falls/Clyburn, and French Mountain), and population size is represented by the number of individuals heard at each station. The salamander data were collected from 13 sites across six time series (2009–2010, 2009–2013, 2009–2016, 2011–2012, 2013–2014, and 2015–2016). Latitude and longitude were also provided in UTM for each site. Habitat consists of temperature in degrees Celsius grouped into three classes: low (<10°C), medium (10–15°C), and high (>15°C). Population size is the number of individuals present at each site. The combinations of data from different time series were performed to assess how the dynamics of the two populations behave over these different time intervals.

The third dataset consists of data on 98 plant species from the Brazilian Cerrado, collected from five surveys. Coordinates were provided in decimal degrees for each site, habitat corresponds to four vegetation types (rock savanna, savanna woodland, savanna, and valley forest), and population size is the number of individuals per plot. Although the surveys were conducted in different years, the interval between the first and second surveys was four years for all of them, allowing comparison of the dynamics of species shared among sites but derived from different data sources. This approach demonstrates the flexibility of the method in operating with data from different collections, which is important in regions where systematic databases are not yet publicly available.

The final database used was a subset of RivFishTIME, a global database that compiles data from monitoring programs and scientific research conducted over decades (Comte et al., 2021). The database contains fish community records collected between 1951 and 2019, spanning multiple river basins and biogeographical areas, allowing for the study of long-term changes in species composition and abundance. The application of RivFishTIME requires consideration of spatial and methodological biases, given the strong concentration of time series in the Palearctic, particularly Europe, the under-representation of tropical realms, and differences in sampling procedures and effort among monitoring programmes. While RivFishTIME is an essential data source for studying long-term changes in freshwater fish communities, results should be interpreted in light of both the data’s poor regional coverage and the database’s methodological variability. For the present study, a subset of the database covering the period 2010–2020 was used. The extracted data included species identity, number of individuals recorded, geographical coordinates, and the water body in which each record was collected.

## 4. Result

The method was able to detected the temporal variation in the three parameters examined, geographical range (*dgr*), habitat specificity (*dhs*), and population size (*dps*), across both single-species and multispecies datasets (Figure 2). Although most estimated changes were relatively small, their temporal patterns revealed differences in the magnitude and direction of change among species and datasets. In particular, the results indicate that temporal variation was not necessarily consistent across parameters, with some species showing changes in population size or habitat specificity while maintaining relatively stable geographical ranges.

**Figure 1.**
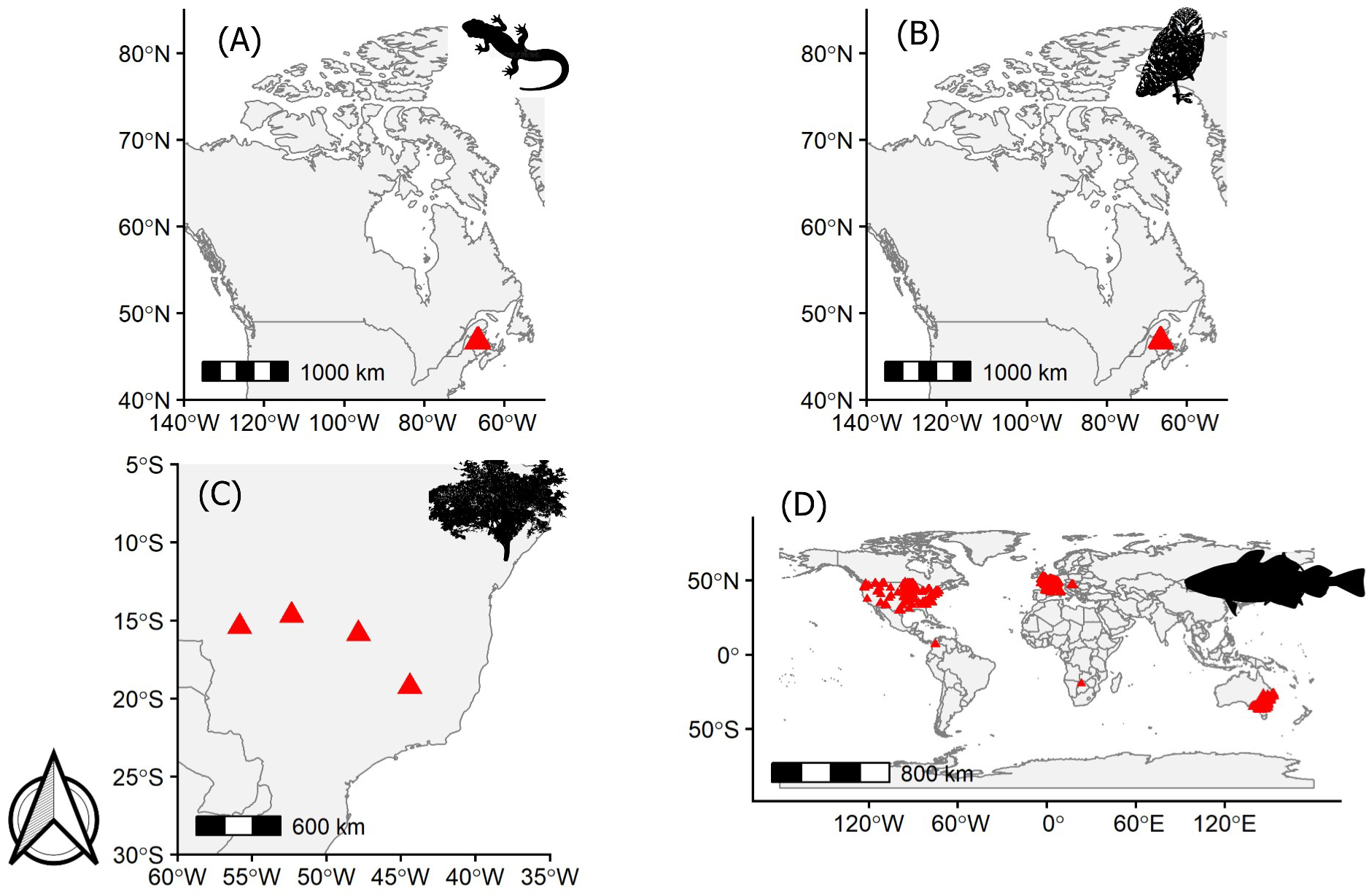
Dataset locations of four different groups. (A) salamander population and (B) owl from Canada, (C) woody community species from Brazil and (D) fish community across the world. Details on data and their references will be made available in a dataset for package-specific testing in the final publication.

**Figure 2.**
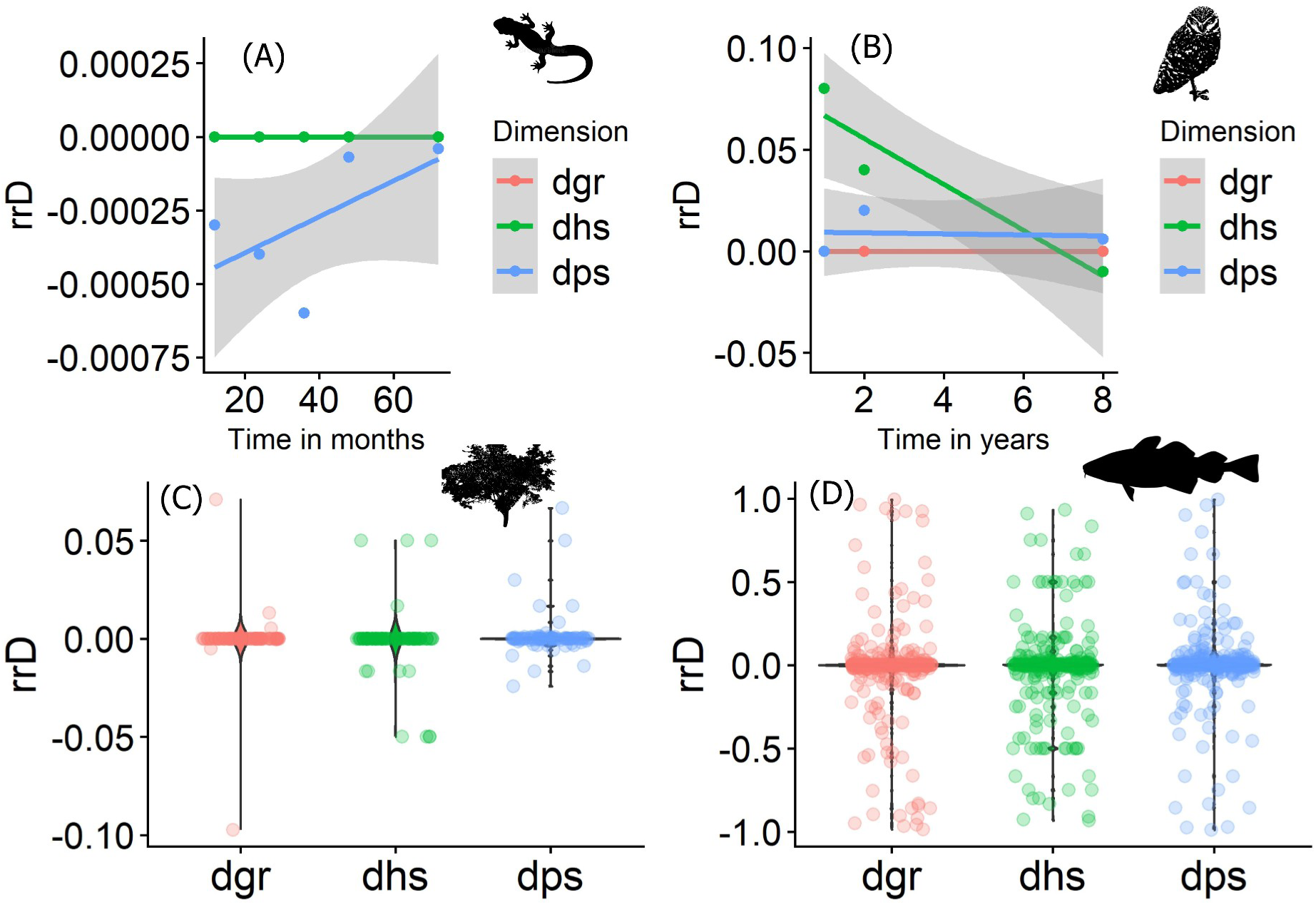
Rarity dynamic (*rrD*) from four different groups: salamander, owl, woody and fish. The metrics were calculated using a new R package developed for this purpose. (A) salamander population and (B) owl from Canada, (C) woody community species from Brazil and (D) fish community across the world.

For the salamander, the temporal patterns were characterised primarily by changes in population size. Population-size estimates were consistently negative across the earlier sampling intervals, reaching their lowest values at approximately 40 months, followed by a progressive increase toward zero at 72 months. This pattern suggests an initial reduction in population size followed by a partial recovery over the longer time interval. In contrast, habitat specificity remained essentially unchanged throughout the study period, with estimates consistently close to zero. Geographical range also remained close to zero, indicating little detectable temporal variation in the spatial extent of the species’ range. Thus, among the three parameters, population size exhibited the clearest temporal signal for the salamander, whereas geographical range and habitat specificity remained comparatively stable.

The owl exhibited a different temporal pattern. Geographical range remained close to zero across the analysed time intervals, indicating little evidence of temporal change in the extent of the species’ geographical distribution. Population size estimates were also relatively small and remained close to zero, with only minor variation among time interval. In contrast, habitat specificity showed a pronounced temporal trend, with positive estimates at the shorter time intervals and progressively lower values over time, becoming slightly negative at the eight-year interval. This pattern indicates a shift from positive to negative changes in habitat specificity over the course of the study. Consequently, the temporal dynamics of the owl were driven primarily by changes in habitat specificity rather than by substantial changes in geographical range or population size

The multispecies dataset of woody plants showed a markedly different pattern from the single-species datasets. Across all three parameters, most species exhibited estimates concentrated around zero, indicating relatively small temporal changes for the majority of species. Nevertheless, positive and negative deviations were observed for geographical range, habitat specificity, and population size, demonstrating variation among species. Geographical range showed particularly evident interspecific variation, including species with comparatively large positive and negative changes. Habitat specificity and population size also exhibited occasional deviations from zero, although most estimates remained tightly clustered around the central value. These results indicate that temporal responses within the woody plant community were heterogeneous, with different species exhibiting increases, decreases, or little detectable change depending on the parameter considered.

The Neotropical fish dataset exhibited the broadest distribution of temporal change estimates among the datasets analysed. For geographical range, habitat specificity, and population size, most species had estimates concentrated around zero, indicating relatively small changes for across species. However, in contrast to the woody plant community, the fish dataset included a much broader range of interspecific responses. Estimates extended across most of the −1 to 1 scale for all three parameters, with species showing both positive and negative changes. This broad distribution indicates substantial heterogeneity in the magnitude and direction of temporal responses among fish species. Despite the strong concentration of estimates near zero, the occurrence of values approaching both extremes of the scale demonstrates that a subset of species experienced considerably larger changes in geographical range, habitat specificity, or population size.

## 5. Discussion

Earlier methods looked at a taxon’s rarity at specific points in space and time (Perrin and Waldren, 2020). In contrast, this approach treats Rabinowitz’s dimensions as state variables that change together in a continuous three-dimensional space. For each dimension, researchers set a boundary between uncommon and common species using an origin point and an empirical threshold (Stenger-Kovács et al., 2025). The taxon’s current position in this space is shown by a directed segment, or versor. Its direction shows the mix across the axes without needing arbitrary weights, and its length, found using the Euclidean norm, gives the overall rarity distance from the origin. The system’s changes over time are described by an ecological vector field. The directional derivatives of this field show both the current velocity and the total movement over time. This way of modelling lets researchers calculate rate changes, movement sizes, and the conditions for ecological balance, which occur when the sum of the directional derivatives is zero. This means the community’s path is stable over time. By focusing on rate vectors and movement speeds in Rabinowitz space, the method shifts the research question. Instead of just asking how rare a species is at one moment, it measures how the traits that make it rare change over time due to regional ecological factors (Belmont et al., 2022; Maciel and Nürk, 2025; Zeni et al., 2025).

The methodological novelty provided here is the change from a static, snapshot-based definition of rarity to a dynamic formulation based on vector calculus and connected state variables. Prior methods assessed taxon rarity at specific spatial and temporal intervals (Perrin and Waldren, 2020); however, this approach reconceptualises Rabinowitz’s dimensions as state variables that progress together within a continuous three-dimensional Cartesian space. An operational border between uncommon and common species categories is established for each dimension by an origin and an empirical cut-off threshold (Rabinowitz, 1981; Stenger-Kovács et al., 2025). The instantaneous position of a taxon in this three-dimensional space is represented by a directed segment or versor whose orientation defines the proportionate combination across axes without imposing arbitrary weighting coefficients, and whose magnitude, computed using the Euclidean norm, measures the total composite rarity distance from the origin. The temporal behaviour of the system is governed by an underlying ecological vector field, whose directional derivatives yield the instantaneous velocity vector and cumulative net displacement over time. This representation allows researchers to calculate rate shifts, displacement magnitudes, and ecological equilibrium conditions in which the sum of directional derivatives vanishes, indicating temporal stability in community trajectories. By focusing on rate vectors and displacement velocities within Rabinowitz space, the method shifts the research question from determining how rare a species is at an isolated moment to quantifying how the traits underlying its rarity change through time under regional ecological drivers (Belmont et al., 2022; Maciel and Nürk, 2025; Zeni et al., 2025).

Using this multidimensional framework with different ecological datasets shows that patterns can vary widely and highlights why it is important to keep track of species-level details. For example, in the case study, salamanders and owls followed different trends over time. Salamanders changed most in population size, while their geographical range and habitat use stayed about the same. Owls, on the other hand, changed most in habitat use, with their population size and range staying more stable. This difference shows that even closely studied groups can respond in unique ways, even in small areas (Lerm et al., 2023; Perrin and Waldren, 2020). When looking at woody plants, the overall community seemed stable, but there was actually a lot of variation between species, especially in their geographical ranges (Maciel and Martins, 2021). So, while the community as a whole might look steady, individual species can be quite unstable, especially those with specific roles (Maciel and Nürk, 2025). The clearest example of this variation was seen in Neotropical stream fish, where some species increased, some decreased, and others stayed about the same. These mixed trends in the same habitat are shaped by past land use and environmental factors that affect common and rare species differently (Lengyel et al., 2025; Mehner et al., 2025; Zeni et al., 2025). In streams and island ecosystems, rare fish and plants often act as important connectors and help maintain diversity, so conservation efforts need to match their unique types of rarity (Stenger-Kovács et al., 2025; Testolin et al., 2024). Comparative evaluation across datasets thus supports the premise that rarity cannot be treated homogeneously and requires systematic dimensional disaggregation to capture community assembly dynamics and species persistence (Forti et al., 2024; Huber et al., 2025).

The implications of this vector-based approach for rarity conservation and species monitoring programs concern the distinction between intrinsic rarity and extinction risk and the identification of changes within individual rarity dimensions. Rarity describes the structural position of a taxon in 3D parameter space, whereas extinction risk depends on the interaction of these parameters with external anthropogenic threats, such as land conversion, fire regimes, and human footprint (Maciel et al., 2023). Monitoring versor velocity and direction along individual axes can also provide an early warning system for conservation programs. As demonstrated in empirical comparative studies, severe demographic crashes can occur within a species’ geographical distribution without immediately reducing its total spatial extent (Zeni et al., 2025). Monitoring frameworks based solely on occurrence extent or spatial occupancy may therefore overlook internal population declines until territorial retreat has occurred (Belmont et al., 2022; Forti et al., 2024). Disaggregating the three dimensions identifies which component of a species’ ecological condition is changing, guiding targeted management interventions: population declines require demographic protection and genetic management, habitat restriction demands strict microhabitat preservation, and range contractions necessitate ecological corridor establishment (Huber et al., 2025; Perrin and Waldren, 2020). This specificity can help direct limited conservation resources toward appropriate management actions while avoiding unnecessary intervention for naturally rare taxa maintaining stable dynamic equilibria (Maciel and Nürk, 2025; Stenger-Kovács et al., 2025). Identifying rare species that serve as functional backbones can also support the preservation of ecological roles during rapid environmental change (Leitão et al., 2016; Mehner et al., 2025). Tracking multidimensional state trajectories thus connects theoretical macroecology with practical conservation decision-making across threatened biological communities (Maciel and Nürk, 2025; Testolin et al., 2024).

Despite these conceptual and analytical advantages, the proposed framework has limitations that point to directions for future research. A primary limitation is the sensitivity of field survey data to imperfect detection and sampling bias, which can produce erroneous rarity classifications or artificial shifts in habitat specificity and population size (Belmont et al., 2022; Mao and Colwell, 2005). In short-term field surveys or studies of cryptic taxa, unobserved species may reflect detection failures rather than true spatial contractions (Belmont et al., 2022; Forti et al., 2024). Temporal sampling scale is a second challenge since time length has significant effects on apparent species abundance distributions and could associate rare species with sporadic or transient species that are caught during brief dispersion pulses (Mehner et al., 2025). Extending time series and deploying repeated multi-interval sampling designs could help distinguish stochastic inter annual variation from sustained, directional trajectories in Rabinowitz space. One way to increase the spatial and temporal precision of state vector coordinates and close geographical data gaps across megadiverse regions is to integrate large-scale citizen science repositories and historical museum collections (Forti et al., 2024; Lerm et al., 2023). Finally, predictive modelling of future rarity trajectories under global climate and land-use scenarios may also be made possible by directly incorporating extrinsic environmental factors, such as climatic anomalies, landscape fragmentation indices, and land-use changes, into the 3D vector field (Maciel et al., 2023; Testolin et al., 2024; Zeni et al., 2025).

This work treats rarity as a dynamic trajectory of linked state variables that change over time, extending Rabinowitz’s (1981) tripartite framework and Maciel’s (2021) synthetic index into a continuous three-dimensional geometric space. Decoupled ecological processes and growing species vulnerabilities that might go undetected by aggregated metrics or unidimensional analyses are revealed by independent temporal tracking along geographical range, habitat specificity, and population size. The technique is used to capture equilibrium states and velocity shifts under different environmental disturbance regimes in freshwater fish, birds, amphibians, and terrestrial plants. This vector-based approach offers a quantitative foundation for differentiating between intrinsic rarity and direct extinction likelihood, as well as between short-term stochastic variation and long-term ecological decline. By enabling dimension-specific management strategies, the approach can support resource allocation in long-term monitoring programs and the maintenance of community functional breadth and ecosystem resilience under environmental change. The framework therefore shifts attention from the static measurement of rarity toward the temporal dynamics of its constituent attributes. As a result, the framework focuses on the temporal dynamics of its component qualities rather than the static measurement of rarity.

## Aknowledgments

This study is part of an ongoing research project initiated in 2020 with support from CNPq (Brazil’s National Council for Scientific and Technological Development), with further funding enabling the research to continue (#200926/2024-1).

